# Myricetin Attenuates LPS-induced Inflammation in RAW 264.7 Macrophages and Mouse Models

**DOI:** 10.1101/299420

**Authors:** Wei Hou, Siyi Hu, Zhenzhong Su, Qi Wang, Guangping Meng, Tingting Guo, Jie Zhang, Peng Gao

## Abstract

**Background:** Myricetin has been demonstrated to inhibit inflammation in a variety of diseases, but little is known about its characters in acute lung injury (ALI). In this study, we aimed to investigate the protective effects of myricetin on inflammation in lipopolysaccharide (LPS)-stimulated RAW 264.7 cells and a LPS-induced lung injury model.

**Methods:** Specifically, we investigated its effects on lung edema and histological damage by lung W/D weight ratio, HE staining and Evans Blue dye. Then macrophage activation was detected by evaluating the TNF-α, IL-6 and IL-1β mRNA and protein iNOS and COX-2. Myricetin was used to detect the impact on the inflammatory responses in LPS-induced RAW264.7 cells with the same manners in mouse model. Finally, NF-κB and MAPK signaling pathways were investigated with Western blot assay in LPS-induced RAW264.7 cells.

**Results:** Myricetin significantly inhibited the production of the pro-inflammatory cytokines *in vitro* and *in vivo*. The *in vivo* experiments showed that pretreatment with Myricetin markedly attenuated the development of pulmonary edema, histological severities and macrophage activation in mice with ALI. The underlying mechanisms were further demonstrated *in vitro* that myricetin exerted an anti-inflammatory effect through suppressing the NF-κB p65 and AKT activation in NF-κB pathway and JNK, p-ERK and p38 in mitogen-activated protein kinases signaling pathway.

**Conclusion:** Myricetin alleviated ALI by inhibiting macrophage activation, and inhibited inflammation *in vitro* and *in vivo*. It may be a potential therapeutic candidate for the prevention of inflammatory diseases.

## Introduction

Acute lung injury (ALI) is defined as an intense and severe inflammatory process in lung, which leads to a considerably high morbidity and mortality in critical patients (Jeyaseelan et al., 2004). ALI is usually considered to be induced by direct causes (e.g. aspiration pneumonia and inhalation injury) and indirect ones (e.g. sepsis, pancreatitis, and blood transfusion) that are related to inflammatory processes including neutrophil accumulation into the alveoli, pulmonary interstitial edema, alveolar epithelium injury, as well as increase of micro-vascular endothelial permeability (Jeyaseelan et al., 2004; Wu et al., 2016).

Pulmonary exposure of lipopolysaccharide (LPS) which is known as endotoxin, is a composition derived from the outer membranes of Gram-negative bacteria, has been reported to trigger macrophages activation and inflammatory cells leakage, especially neutrophils (Idell, 2003; Jeyaseelan et al., 2004). Previous studies demonstrated that LPS activated inflammatory signaling pathway nuclear factor-kappa B (NF-κB) and mitogen-activated protein kinase (MAPK) (Hayden and Ghosh, 2008; Schuliga, 2015). Accumulating evidence supports that the activation of the above two signaling pathway in lung tissue is up-regulated in mice processed LPS-induced ALI (Lv et al., 2017; Wu et al., 2016), while inhibitors of these signaling pathway not only suppressed the influx of neutrophils but also alleviated lung edema (Everhart et al., 2006; Lin et al., 2015; Santos et al., 2018).

Myricetin (3,3′,4′,5,5′,7-hexahydroxyflavone) is an abundant flavonoid compound extracted from natural ingredients such as berries, herbs and vegetables (Salvamani et al., 2014). It contributes to the prevention and/or delay of various pathological processes including cancer, neurodegenerative diseases, and inflammatory disorders (Choi et al., 2017; Domitrovic et al., 2015). However, little is concerned about its preventive response on LPS-induced lung injury. In the present study, we investigated the anti-inflammatory influences of myricetin in RAW 264.7 macrophages and the protective activity in LPS-induced acute lung injury mouse model. The purpose of this research was to estimate the prophylaxis function of myrecitin on LPS-induced lung injury *in vitro* and *in vivo*, and elucidate the underlying mechanism.

## Materials and methods

### Reagent

Myricetin with a purity of 99% was purchased from PufeiDe Bio-Technology (Chengdu, China). Primary antibodies (ERK1/2, p-ERK1/2, AKT, p-AKT, P38, p-P38, JNK1/2, p-JNK1/2, NF-κB-p65, and NF-κB-p-p65) were procued from Cell Signaling Technology (Boston, USA). Primary antibodies COX-2 and iNOS were obtained from Abcam (Cambridge, UK), Primary antibody β-actin and HRP-conjugated anti-mouse or anti-rabbit secondary antibodies were acquired from Bosterbio (USA)

### Animals

BALB/C mice in both gender (8–10 weeks of age, 25–30 g weight) were purchased from the Experimental Animals Center of Norman Bethune Medical College of Jilin University (Changchun, PRC). The mice were separated by sex gender and maintained under a same pathogen-free condition for 1 week before the start of experiment. Animals were maintained at 22-24 °C with an artificial 12h/12h day/night cycle. All the animals was provided with adequate food and drink.

### Animal treatment

One hundred and twenty BALB/C mice were distributed into five treatment groups: (i) sham group (n=24), treated with 20μL PBS; (ii) LPS group, subject to LPS (50 μg) dissolved in 20μL PBS via intranasal administration to induce ALI; (iii) LPS+Myr groups (n=24 per group), subject to LPS and 2.5, 5and 10 mg/kg myricetin, respectively. Myricetin was injected intraperitoneally as a pretreatment for 1h. Mice of the sham and LPS groups took in an equivalent of vehicle instead. 12hrs after LPS treatment, the mice were euthanized by Pentobarbital Sodium injection (1%). The animal treatment was conducted within the prescribed limits of Animal Welfare Committee of Jilin University (File No. 2015047).

### Lung wet-dry weight (W/D) ratio

About 12 hrs after intranasal LPS instillment, lung tissues from 4 mice per group not subjected to BALF (Bronchoalveolar Lavage Fluid) were excised. The surface liquid of lung section was drained with clean filter paper carefully and then record as the wet weight. Stored them in a thermostatic drier at constant 80°C for 48 hrs then the dry weight were obtained. The W/D ratio was calculated to evaluate the moisture content of lung tissue.

### MPO activity measurement

MPO activity was measured to monitor the parenchymal infiltration of macrophages and neutrophils. In this study MPO activity was assayed with a commercial test kit (Jiancheng bioengineering institute, A044, Nanjing, PRC). Briefly, mice were killed 12 hrs after LPS treatment by diethyl ether deep anesthesia. Lung tissues were collected and weighted and then homogenized with reagent in the test kit followed by centrifuged according to the specification. The supernatant of cell-free extracts were collected afterwards and stored in -20°C before usage. Terminate the reaction and measure the absobance at 450 nm in a microplate reader.

### HE staining

Histopathologic evaluation was processed to the mice that not undergo the collection of BALF. The collected lung tissue was soaked in 10% neutral buffered formalin for 24hrs to immobilize the structure at first, embedded in paraffin subsequently, sliced at the thickness of 5μm and ultimately stained with hematoxylin and eosin (H&E).

### Lung barrier permeability

The mice were injected with 2% EBD (100 mg/kg, Sigma-Aldrich) via the tail vein 30 min before euthanization. After euthanization, the lung tissues were removed *en bloc* and homogenized, and the homogenate was incubated at 37 °C for 24 hrs. Then blue dye was performed at 620 nm by using a microplate reader.

### Cell culture and treatment

RAW264.7 mouse macrophage were acquired from the China Cell Line Bank (Beijing, PRC). Cells were resuscitated and cultured in DMEM complete medium containing 10% fetal bovine serum (Gibco, USA) at 37°C in thermostatic incubator with 5% CO2. After revive from cryopreservation and continuous cell culture, cells in logarithmic growth phase were placed in 6-well plate and cultured overnight. After that change the medium to serum-free medium hence the cells were starved for 4hrs. Then the cells were pretreated with different gradient concentrations (12.5μM,25μM,50μM and 100μM) of myricetin about 1 hr, afterwards incubating with LPS (1μg/mL) for 12 hrs. Subsequently, the serum-free solution was extracted and centrifuged at 10000g maintained 4°C for 15min to collect the supernatant. TNF-α, IL-6 and IL-1β secreted extracellular were measured using BioLegend ELISA kit (San Diego, USA), follow the producer’s specification.

### MTT assay for cell viability

RAW 264.7 cell suspension were purfuse into 96-well plates containing 100 μL of DMEM at a density of 5×10^5^ cells/mL. The mixture was incubated in an thermostatic chamber at 37°C in 5% CO_2_ for 1 hr. Then the cells were preprocessed with 50 mL of myricetin (12.5–100μM), 1 hr later, the stimulation with 50 mL LPS solution(1 mg/mL) was added to each well. After18 hrs, 20μL MTT (5 mg/mL) was added per well, and undergo further incubation for another 4 hrs. The supernatant was discarded and then DMSO wre added 150 μL/well to lyse the colored crystal. Optical density of the solution was detected at 570 nm absorbance on a microplate reader.

### Quantitative Real-Time PCR

The qRT-PCR was carried out according to our previous description (Lv et al., 2016). Briefly, total RNA of the RAW264.7 cells was extracted with Tri-reagant (sigma-aldrich, USA) follow the manufacturer’s instructions, and the cDNA was generated though a commercial RT-PCR Kit (GeneCopoeia, Maryland, USA), and then analyzed with Takara SYBR Green RT-PCR Kit (Takara, Kyoto, Japan), and every sample was repeated triple times. Sequence of primers is displayed in Table 1. β-actin served as the internal standard.

**Table 1.**
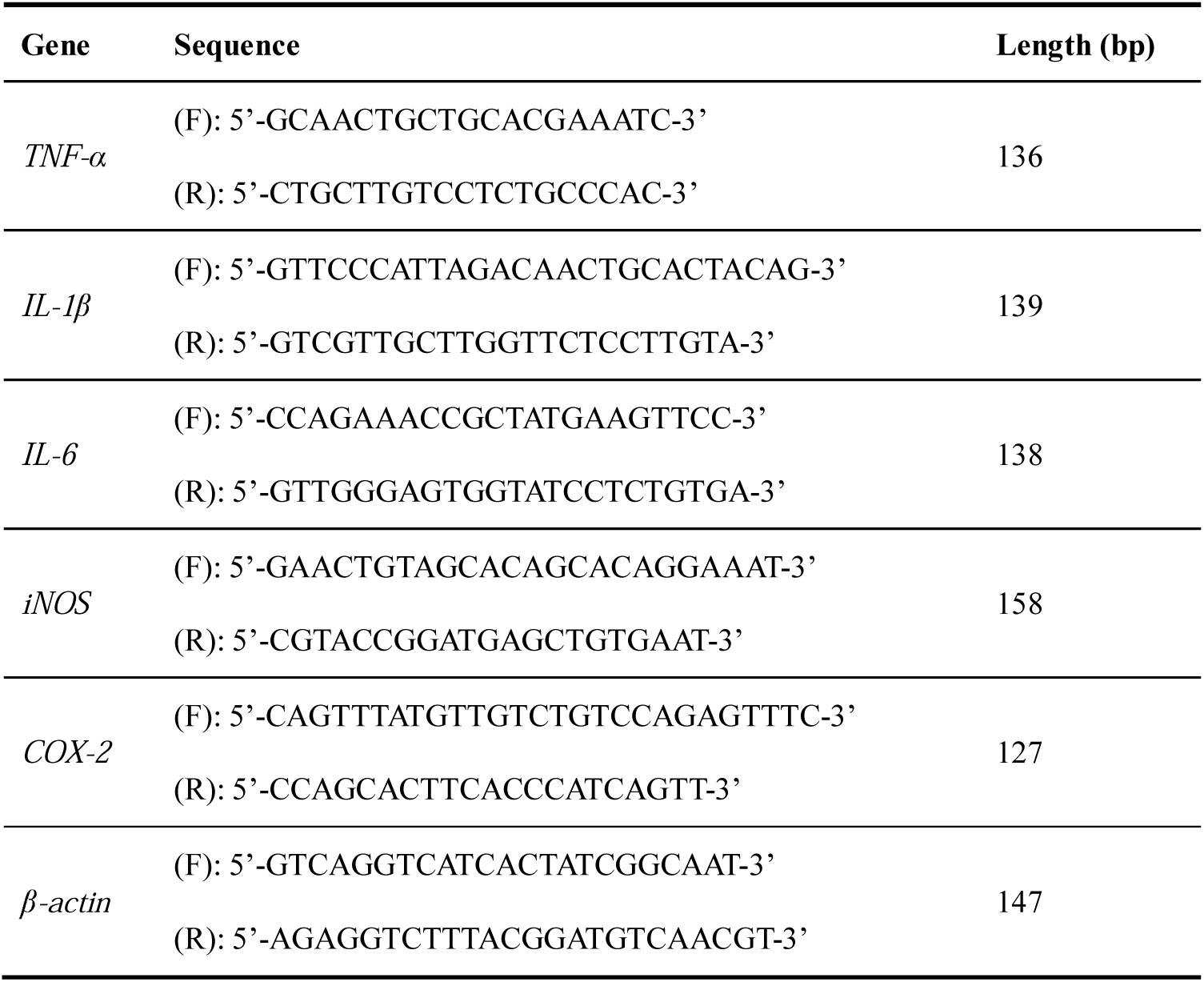
The primer sequences of *TNF-*α, *IL-1*β, *IL-6*, *iNOS*, *COX-2* and β*-actin* (mice)

### Statistics

The results are displayed as mean ± standard deviation (SD). Data were analyzed with statistical software SPSS 19.0 (SPSS Inc, USA). The dissimilarity among groups were analyzed by one-way analysis of variance (ANOVA). All bar charts were accomplished with GraphPad Prism (v7; GraphPad Software, USA). *P*<0.05 was regarded as statistically significant.

## Results

### Myricetin reduced lung histopathological changes induced by LPS

As shown in Fig.1, the structure of lung parenchyma was almost intact in the sham group. The structure in LPS group displayed evident pathologic lesions, principally alveolar wall incrassation and alveolar disarrangement. In the LPS+Myr groups, histopathologic damage were obviously alleviate by myricetin of all concentration(Fig. 1).

**Figure 1.**
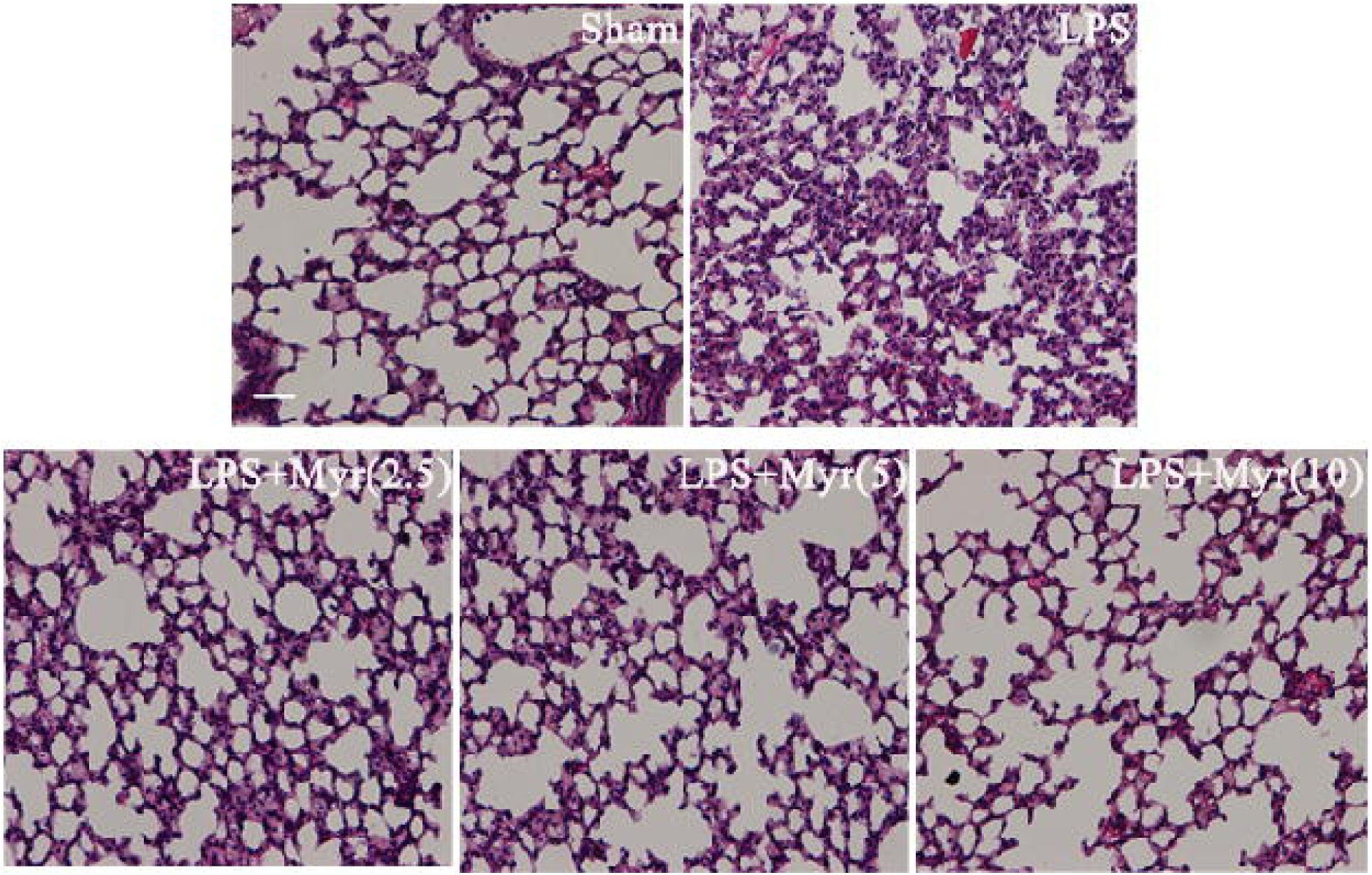
Effects of myricetin on histological changes in lung tissues from mice with lipopolysaccharide (LPS)-induced acute lung injury (ALI) All mice were randomly divided into five groups and each groups contain 12 mice: the sham-operated group; LPS treatment group; LPS and myricetin (2.5, 5 and 10 mg/kg) treatment group. Myricetin was dissolved in sterile saline containing 5% Tween 80 (v/v). Myricetin (2.5, 5 or 10mg/kg) was intraperitoneal injected 1 hr prior to administration of LPS. LPS (50 μg) was dissolved in 20μL PBS and then administrated intranasally to induce acute lung injury Lungs from each experimental group were processed for histological evaluation at 6 h after LPS challenge respectively. Representative histological section of the lungs was stained by hematoxylin and eosin (HE staining, magnification 100, scale bars = 200μm).

### Myricetin alleviated LPS-induced lung barrier dysfunction

LPS-induced ALI was manifested as pulmonary edema caused by lung barrier dysfunction due to the change of pulmonary vascular permeability principally. Such process usually accompanied by inflammatory cell infiltration in alveoli. In this study, we assessed lung barrier dysfunction by Evans blue dye and pulmonary edema though measuring lung W/D ratio, respectively. LPS induced lung vascular leakage and myricetin inhibited LPS-induced increase of permeability as evaluated by measure Evans blue extravasation into the lung tissue (Fig. 2A and 2B). Lung W/D weight ratio was remarkably up-regulated since LPS administration and pre-treatment with myricetin decreased the ratio (Fig. 2.C). MPO activity in lung parenchyma showed significant increase following LPS dispose in comparison with those of the sham group. Meanwhile, myricetin pretreatment with different concentration(1, 2 or 4 mg/kg) could efficaciously decrease the MPO activity of myricetin treateded mice compared with the LPS group (P<0.01, Fig. 2D). Therefore, myricetin inhibited the activiation of inflammatory cells.

**Figure 2.**
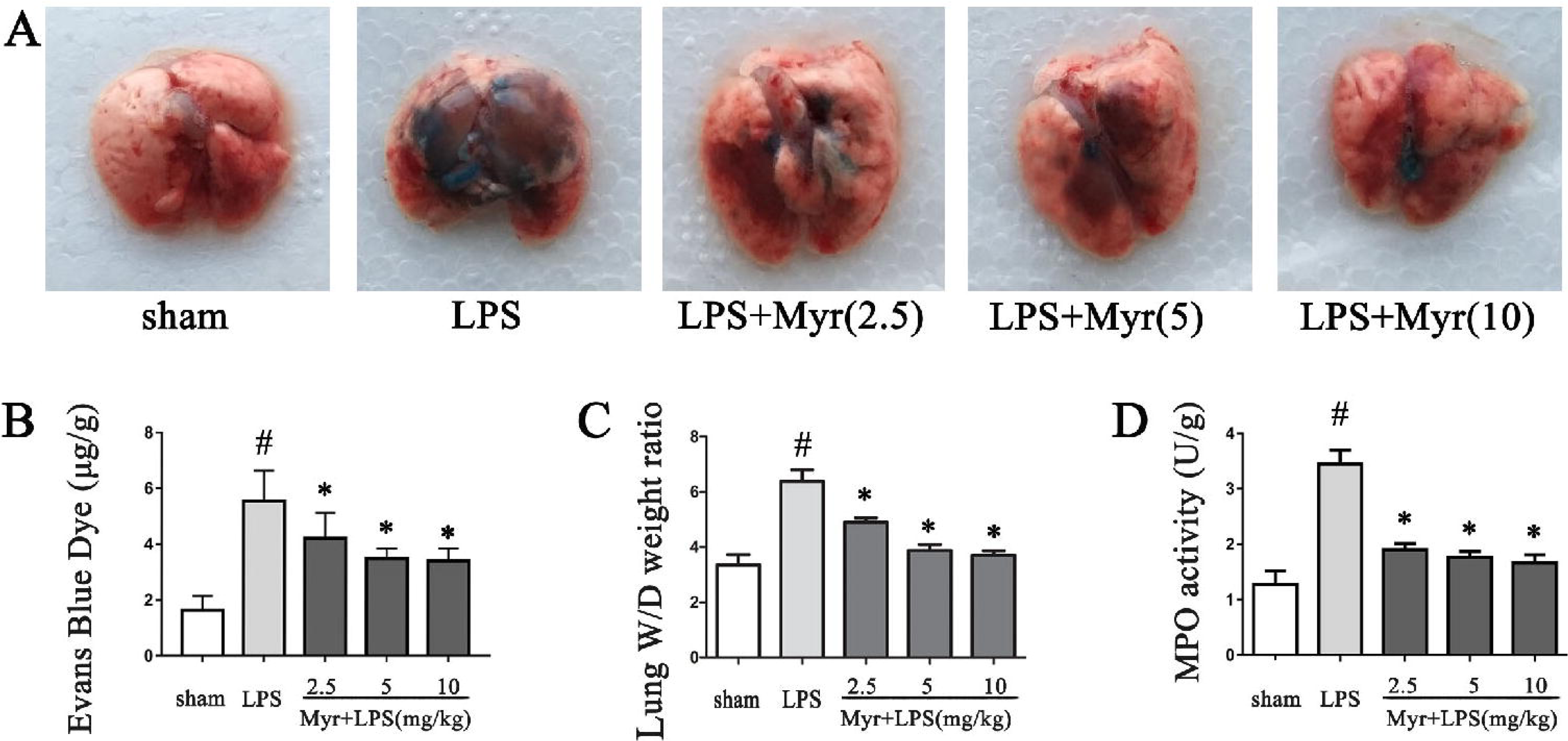
Myricetin alleviated LPS-induced lung barrier dysfunction and edema. (A) Analysis of LPS-induced lung injury and barrier dysfunction. The mice were intravenously injected with Evans blue dye (50 mg/kg) 1hr before termination. An increase was noticed in lung vascular permeability based on the accumulation of Evans blue dye in the lung tissue. (B) Bar graph indicated the quantitative analysis of Evans blue dye extracted from tissue sample. (C) The lung W/D weight ratio was determined at 12 hrs after LPS challenge. (D) MPO activity in lung tissues was measured at 12 hrs after LPS challenge. Similar results were obtained from three independent experiments. All data are presented as means ± SD. #*P* < 0.05 compared to the sham group, \**P* < 0.05 compared to the LPS group.

### Myrecitn decreased the production of inflammation cytokines in BALF and lung tissues

Treatment of LPS induced significant elevation of TNF-α, IL-6 and IL-1β in BALF in comparison with those in control group, while myricetin pre-treatment could induce significant decrease of these factors (Fig. 3A-3C). Expression of TNF-α, IL-6 and IL-1β mRNA by qRT-PCR indicated that myricetin could down-regulate the mRNA expression in comparison with those of the LPS group (Fig. 3D-3F).

**Figure 3.**
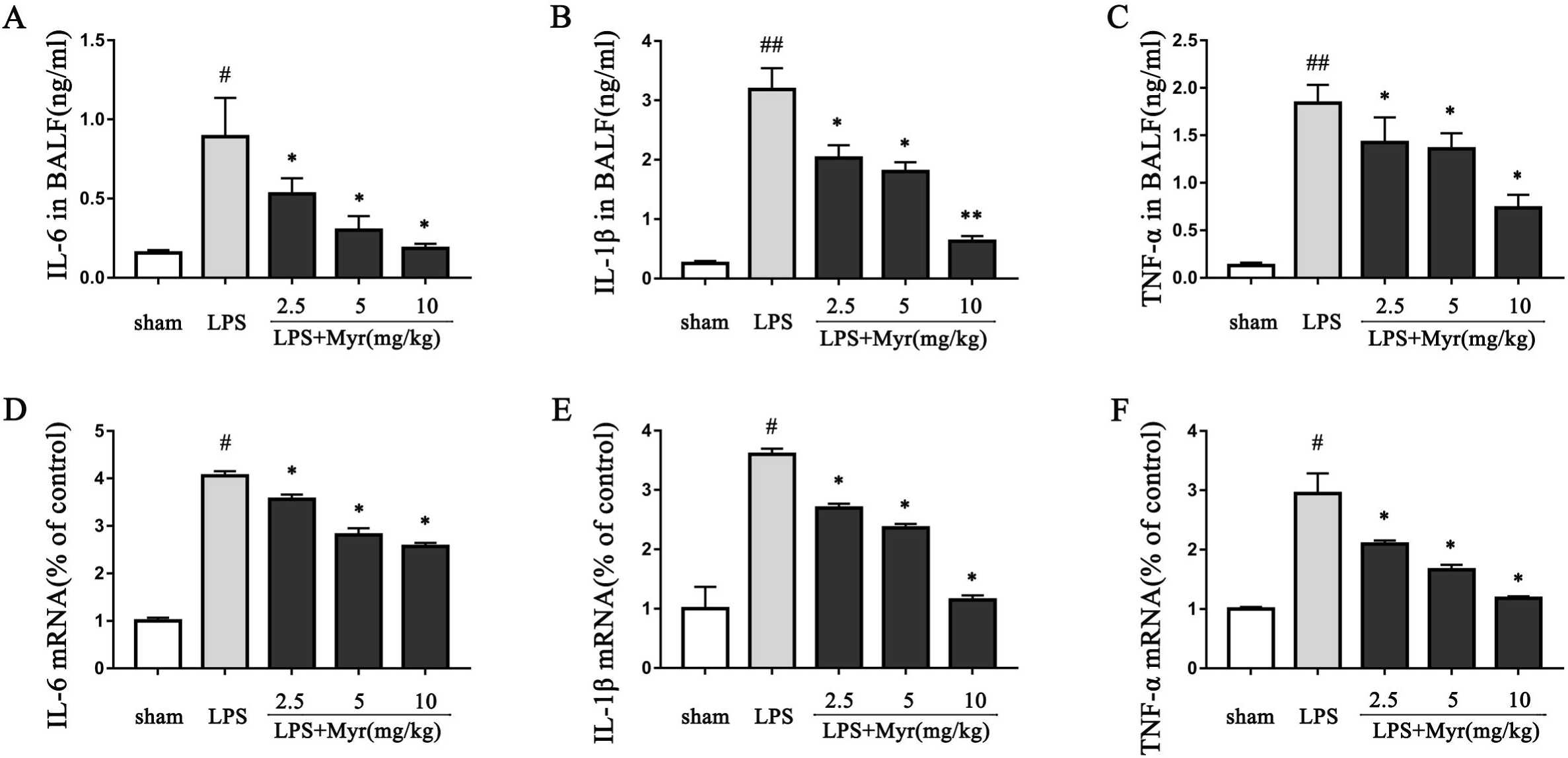
Myriceitn inhibited the activation of inflammation cells in an LPS-induced ALI mouse model. (A-C) The level of cytokines in BALF (*n* = 4 per group). (D-F) The mRNA expression of IL-6, IL-1β and TNF-α in lung tissues (*n* = 4 per group). Data were show as means ± SD. ##*P* < 0.01 compared to the sham group, \**P* < 0.05 and \*\**P* < 0.01 compared to the LPS group.

### Effect of myricetin on cell viability and myricetin restrained the inflammatory response in RAW264.7 cells induced by LPS

MTT assay was proceeded to estimate cytotoxicity of myricetin with several concentrations (12.5mM-100mM). Myricetin showed no cytotoxicity against RAW264.7 under a concentration of 25mM (Fig. 4A). On this basis, 12.5mM and 25mM were selected for the *in vitro* experiments. The effects of myricetin on the expression of iNOS, COX-2, TNF-α, IL-1β, and IL-6 in genetic level were detected by qRT-PCR. Real-time PCR indicated that myricetin reduced the mRNA expression of pro-inflammatory factors TNF-α, IL-6, IL-1β, COX-2 and iNOS induced by LPS(Fig. 4B-4F). Western blot indicated that myricetin significantly reduced the expression of COX-2 and iNOS in both high and low concentration in protein level (Fig. 4G-4I). Furthermore, ELISA result demonstrated that myricetin reduced the generation of TNF-α and IL-6 in LPS-induced RAW264.7 cells (Fig. 5A-5B).

**Figure 4.**
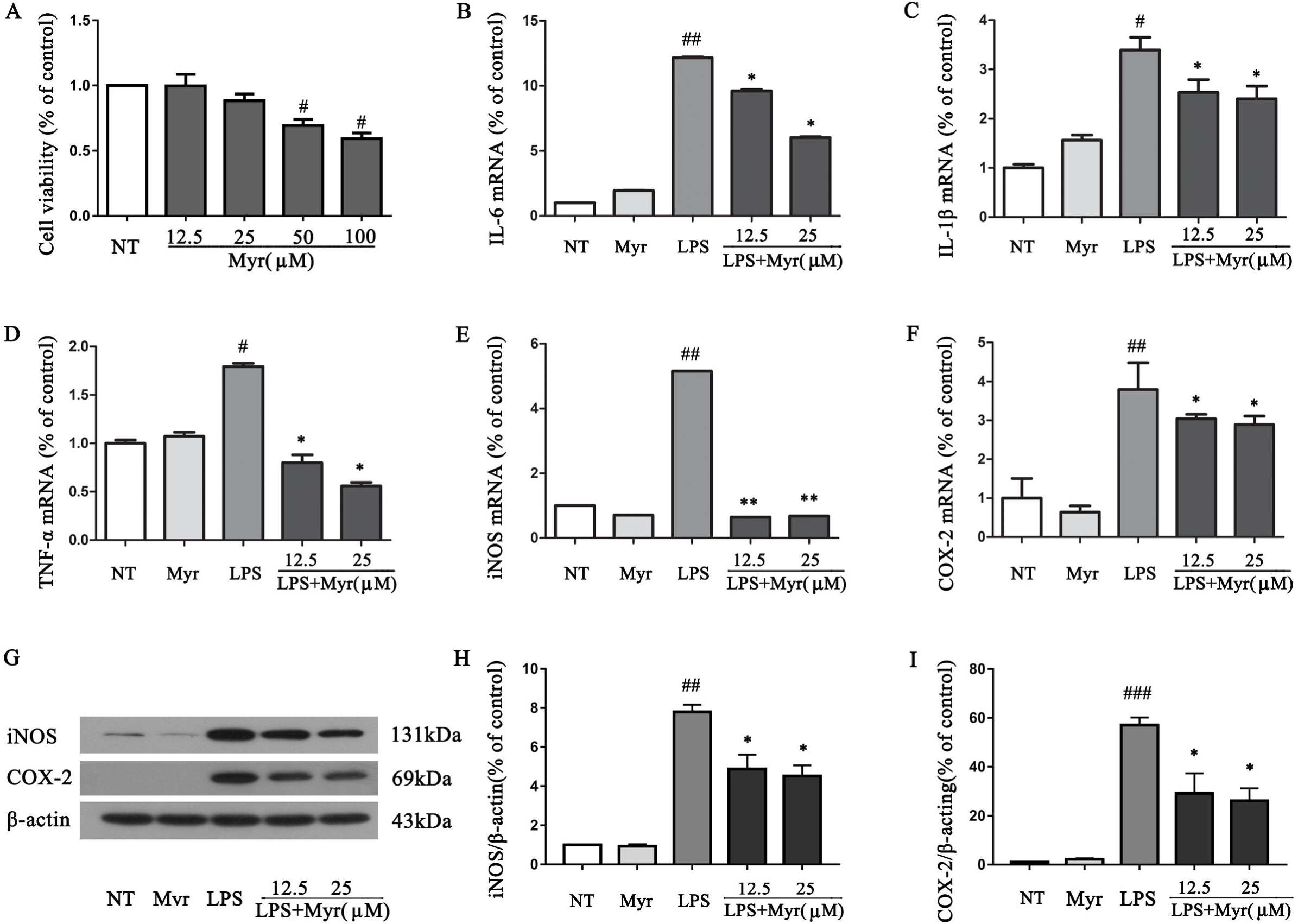
Myricetin suppressed the LPS-induced inflammatory response in RAW264.7 cells. (A) Effects of myricetin on RAW264.7 cell viability (*n* = 5). (B-F) The RAW264.7 cells were stimulated with LPS for 4 hrs after pretreating with myricein for 1 hr, and then cells were collected with Trizol. The mRNA expression of TNF-α, IL-6, IL-1β, COX-2 and iNOS was detected by qRT-PCR (*n* = 3). The protein levels of COX-2 and iNOS was analyzed by western (G-I) (*n* = 3). Data were shown as means ± SD. ###*P* < 0.001 compared to the no-treatment group (NT), \**P* < 0.05, \*\**P* < 0.01 and \*\*\**P* < 0.001 compared to the LPS group.

**Figure 5.**
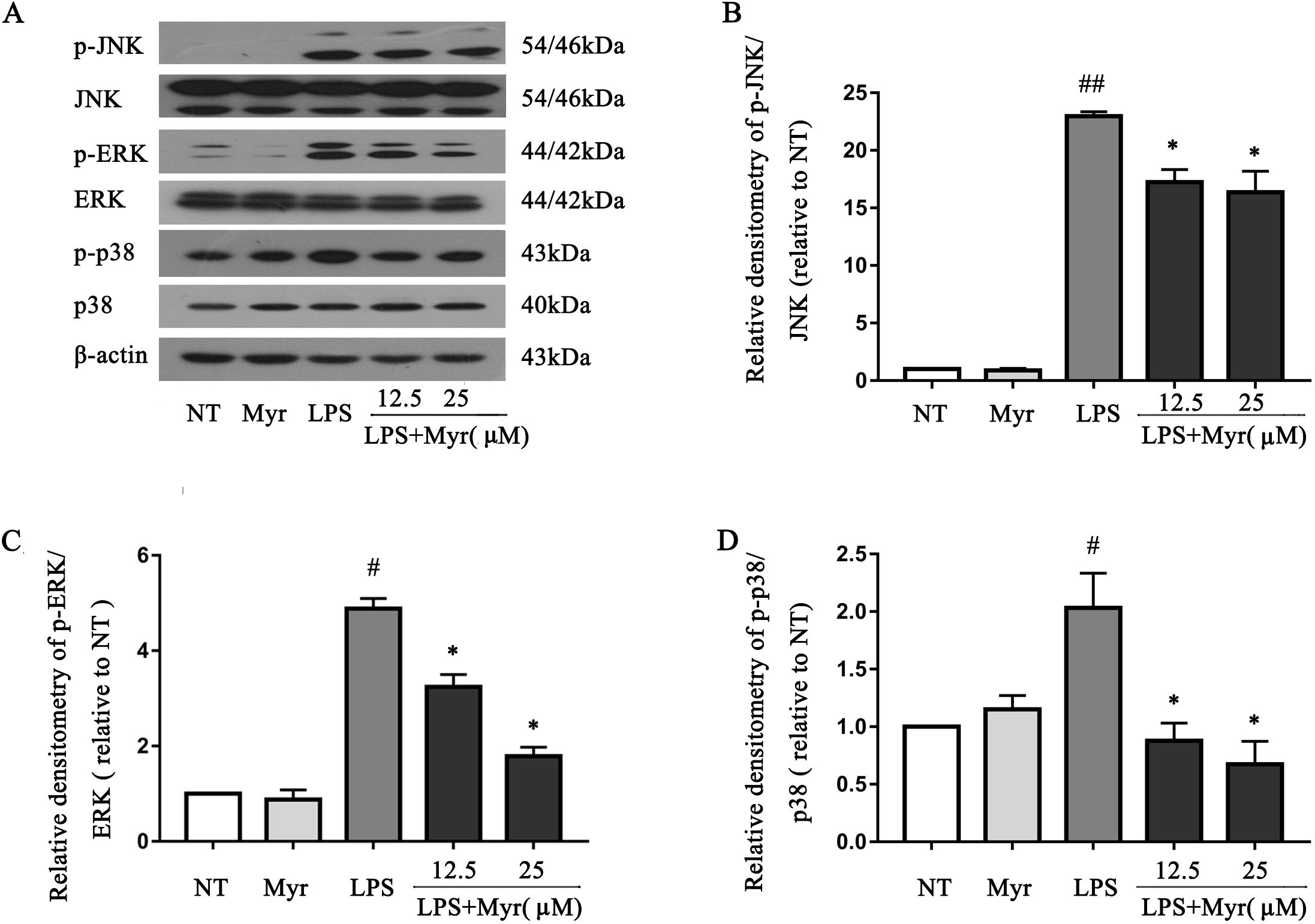
Myricetin inhibited the the secretion of cytokines. RAW264.7 cells were stimulated with LPS for 12 hrs after pretreating with myricetin for 1 hr, and then the supernatant was collected. The protein levels of TNF-α, IL-6 and was analyzed by ELISA (A-B) (*n* = 3). Data were shown as means ± SD. #*P* < 0.05 compared to the no-treatment group (NT), \**P* < 0.05, compared to the LPS group.

### Myricetin modulated the phosphorylation of the p38/MAPKs and p65/AKT signaling pathways in LPS-induced RAW264.7 cells

Phosphorylation of critical factors in these signalling pathways could activate pro-inflammatory reaction in LPS-induced RAW264.7 cells. Inhibition of these factor could suppress the process. In this study, myricetin remarkably associated with phosphorylation of signaling molecule p38, ERK and JNK (Fig. 6) of MAPK signaling in LPS-induced cells. Some key factors in signaling pathway AKT were measured, which indicated that myricetin also explicitly related to the phosphorylation of p65 and AKT (Fig. 7).

**Figure 6.**
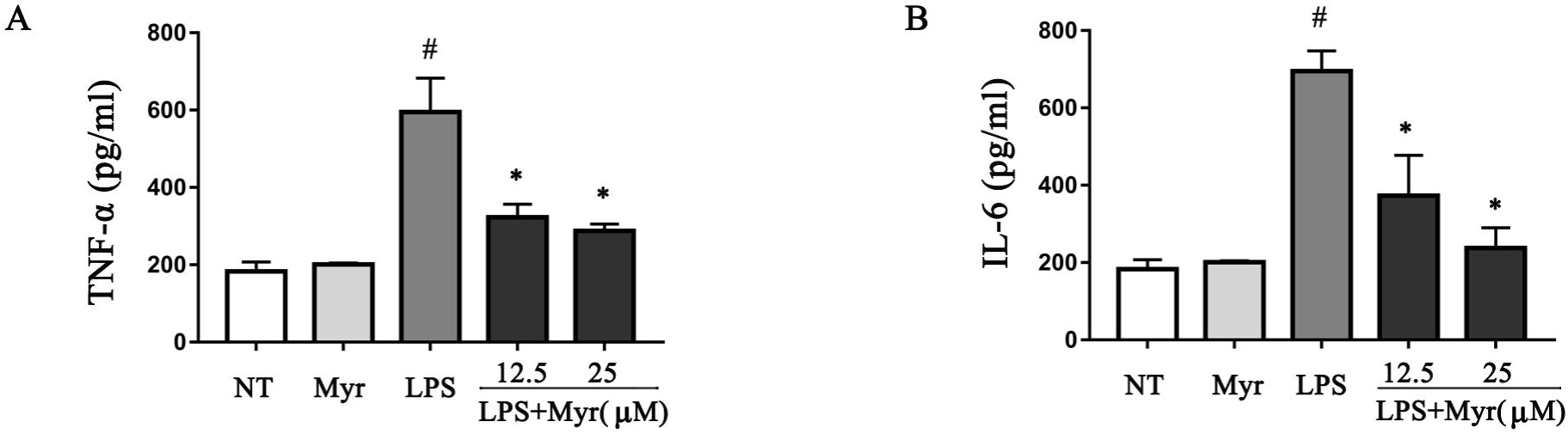
Myricetin inhibited the phosphorylation of JNK, p-ERK and p38. (A) Western blot analysis for the phosphorylation of JNK, ERK and p38 (*n* = 3). Cells were pretreated with various concentrations of myricetin (12.5 and 25mM) for 1 hr, followed by stimulation with LPS for 1 hr before harvested. Western blot analysis was performed using antibodies against phospho-or total forms of JNK, ERK and p38. (B-D) Quantification of western blot data ofp-JNK, p-ERK and p-p38. Data are shown as means ± SD. #*P* < 0.05 and ##*P* < 0.01 compared to the NT group, \**P* < 0.05 and compared to the LPS group.

**Figure 7.**
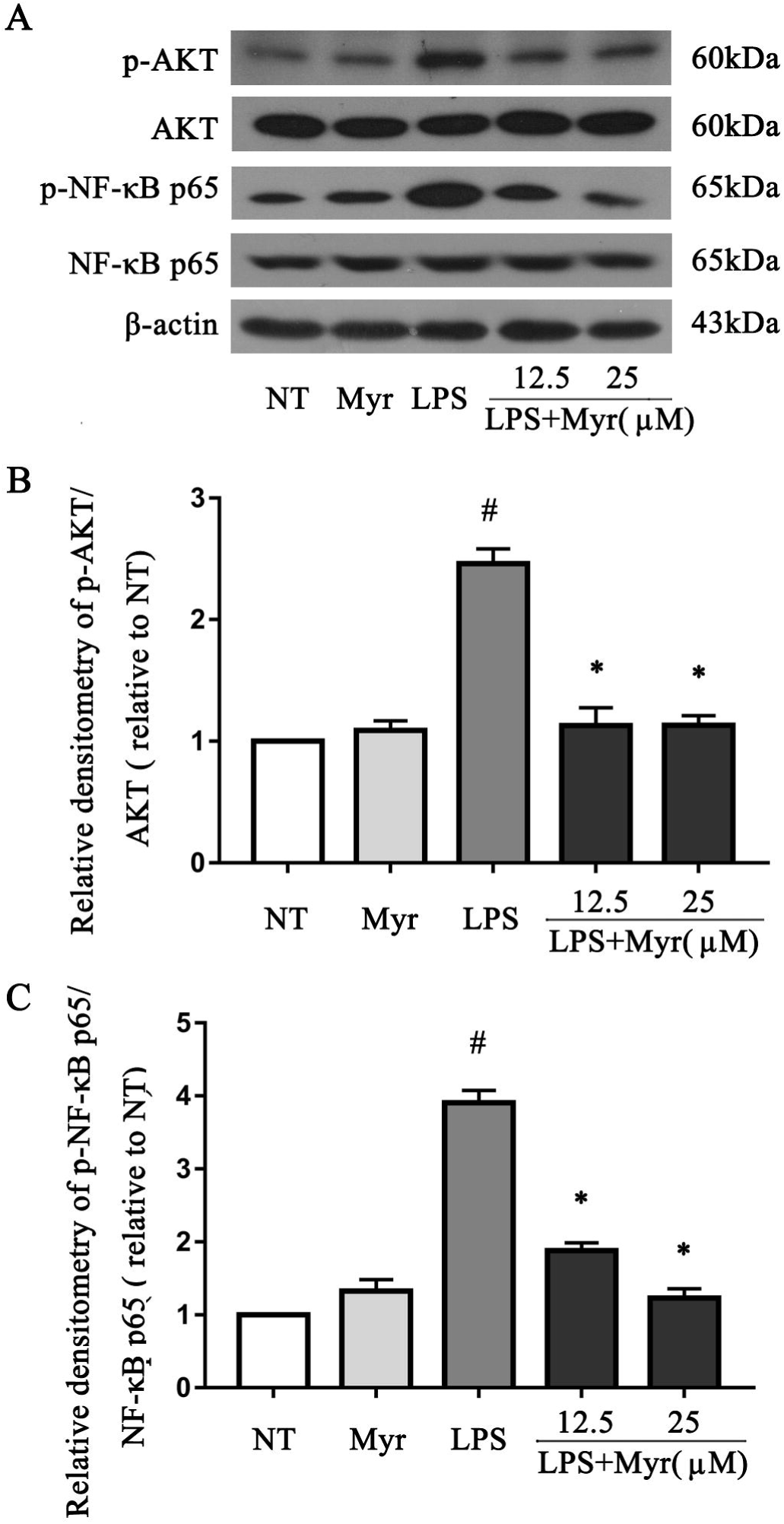
Myricetin inhibited the phosphorylation of AKT and NF-κB p65. (A) Western blot analysis for the phosphorylation of AKT and NF-κB p65 (*n* = 3). Cells were pretreated with various concentrations of myricetin (12.5 and 25mM) for 1 hr, followed by stimulation with LPS for 1 hr before harvested. Western blot analysis was performed using antibodies against phospho-or total forms of AKT and NF-κB p65. (B, C) The relative densitometry of p-AKT and p-NF-κB p65 (relative to normal). Data were shown as means ± SD. #*P* < 0.05 compared to NT, * *P*< 0.05 compared to the LPS group.

## Discussion

ALI is an intense respiratory failure syndrome characterized by uncontrolled inflammatory responses generally induced by various pro-inflammatory cytokines which results in pulmonary edema and damaged alveolar structure (Lv et al., 2017; Xie et al., 2014). LPS is widely used to induce acute inflammation in animal models for the preclinical evaluation of anti-inflammatory drug candidates (Ortiz-Diaz et al., 2013; Shu et al., 2014; Wu et al., 2016). Intranasal injection of LPS is a classical method to establish mouse models of ALI (Zhang et al., 2011), and the symptoms appears in mice with ALI were quite similar to those of human (Proudfoot et al., 2011). Since ALI is a severe clinical syndrome with high mortality and no effective drug is available yet, the prevention of ALI is a vital therapeutic goal (Rubenfeld and Herridge, 2007). In this paper, we measured the anti-inflammation effects of myricetin in *in vitro* and *in vivo*.

Myricetin is a natural flavonol commonly contained in vegetables, fruits, tea and red wine that have been reported to participate in diverse biological effects with various mechanisms (Kang et al., 2010). The estimated intake of myricetin was 0.98 mg/day for female and 1.1 mg/day for male, respectively (Lin et al., 2006). Recent studies revealed that myricetin inhibited cyclooxygenase (COX-2) expression and prostaglandin E_2_ by suppressing NF-κB transcription activation (Lee et al., 2007). In addition, myricetin could also inhibit the nuclear translocation of p65, degradation of IκBα, and cellular apoptosis in LPS-induced cardiac injury (Zhang et al., 2017). Meanwhile, several studies confirmed that NF-κB and MAPKs could mediate inflammatory responses in lung (Santos et al., 2018; Wu et al., 2016). On this basis, we think it was a possible candidate for treating ALI.

Accumulated experiments proved that over-activated inflammatory cells in alveoli mediated the damages of epithelium and vascular endothelium which resulted in barrier dysfunction and the increase in vascular permeability as well as pulmonary edema finally (Bhattacharya and Matthay, 2013; Kim et al., 2017). The lung epithelium barrier integrity of the mice was tested according to the Evans blue dye extravasation assay to reveal the alveolar epithelial completeness. Our data showed that lung W/D weight ratio was assessed as an index of pulmonary parenchyma edema and an indicator of vascular leakage. Myricetin reduced the W/D weight ratio and the accumulation of Evans blue dye in LPS-induce lung injury mice which indicated that myricetin alleviated the vascular and epithelium damage caused by LPS. An imbalance of inflammatory cytokines principally TNF-α,IL-6, IL-1β, iNOS and COX-2 was observed in BALF in LPS-induced lung injury mouse. MPO, the primary marker of neutrophils infiltration in tissues, was also inhibited by myricetin. Pro-inflammatory cytokines initiate an inflammatory cascade and promote the migration of neutrophils into the alveoli. In this present study, myricetin obviously decreased the production of these cytokines both *in vitro* and *in vivo* (Domitrovic et al., 2015; Jung et al., 2017; Jung et al., 2008; Walker et al., 2000).

Macrophage activation induced by endotoxin triggers overproduction of inflammatory cytokines (e.g. IL-6 and TNF-α) that are closely related to the pathogenesis of ALI (Lee et al., 2016). Hence, RAW264.7 macrophage cell line was cultured to imitate the inflammation in alveoli (Chowdhury et al., 2017). ELISA result demonstrated that LPS markedly up-regulated the levels of TNF-α and IL-6 in the culture medium. No significant differences were observed in IL-1β level in the supernatant of the cell culture. Real-time PCR indicated that myricetin dramatically inhibited LPS-induced inflammation by reducing the production of COX-2, iNOS, TNF-α, IL-6 and IL-1β in genetic level. Nevertheless, the underlying mechanism of myricetin suppressed inflammation is still unknown. NF-κB is considered the representative proinflammatory signaling factor based on its character in the expression of genes including cytokines, chemokines, and adhesion molecules (Lawrence, 2009). NF-κB is activated by Toll-like receptors (TLRs) during bacterial and/or viral infection (Hayden and Ghosh, 2008; Sohn et al., 2005). The inhibition of NF-κB/AKT and p38/MAPK signaling could suppress the expression of pro-inflammatory mediators. Hence, to lucubrate possible mechanism of myricetin inhibited inflammatory response induced by LPS in RAW264.7, further measurement of the protein level of NF-κB/AKT and p38/MAPK signaling pathways were performed. The data showed the protective effects of myricetin were associated with inhibition of both NF-κB/AKT and p38/MAPK signaling pathways.

## Conclusions

In summary, myricetin offers a protective role against LPS-induced ALI. High dose of myricetin markedly ameliorated pulmonary edema and inflammation caused by LPS intranasal instillation in mouse model. In future, we will create a new model with primary cells to provide more solid evidence about the ant inflammatory effect of myricetin.

## List of abbreviations

ALI: Acute lung injury
LPS: lipopolysaccharide
NF-Kb: nuclear factor-kappa B
MAPK: mitogen-activated protein kinase
COX-2: cyclooxygenase
TLRs: Toll-like receptors
H&E: hematoxylin and eosin
SD: standard deviation
ANNOVA: one-way analysis of variance

## Acknowledgments

This work was supported by National Natural Science Foundation of China (81672297).

## Authors’ contributions

HW wrote the manuscript; ZJ and GP revised the manuscript; HSY, SZZ, WQ did the data analysis; MGP, GTT did the data collection.

